# Segment 2 from influenza A(H1N1)pdm09 viruses confers temperature sensitive HA yield on candidate vaccine virus growth in eggs that is complemented by PB2 701D

**DOI:** 10.1101/541235

**Authors:** Saira Hussain, Matthew L. Turnbull, Rute M. Pinto, John W. McCauley, Othmar G. Engelhardt, Paul Digard

## Abstract

Candidate vaccine viruses (CVVs) for seasonal influenza A virus are made by reassortment of the antigenic virus with a high-yielding egg-adapted strain, typically A/Puerto Rico/8/34 (PR8). Many 2009 H1N1 pandemic (pdm09) high-growth reassortants (HGRs) selected by this process contain pdm09 segment 2 in addition to the antigenic genes. To investigate this, we made CVV mimics by reverse genetics (RG) that were either 6:2 or 5:3 reassortants between PR8 and two pdm09 strains, A/California/7/2009 (Cal7) and A/England/195/2009, differing in the source of segment 2. The 5:3 viruses replicated better in MDCK-SIAT1 cells than the 6:2 viruses, but the 6:2 CVVs gave higher HA antigen yields from eggs. This unexpected phenomenon reflected temperature sensitivity conferred by pdm09 segment 2, as HA yields from eggs for the 5:3 viruses improved substantially when viruses were grown at 35°C compared with 37.5°C, whereas 6:2 virus yield did not. Authentic 5:3 pdm09 HGRs, X-179A and X-181, were not markedly temperature-sensitive however, despite their PB1 sequences being identical to that of Cal7, suggestive of compensatory mutations elsewhere in the genome. Sequence comparisons of the PR8-derived backbone genes identified single changes in PB2 and NP, 5 in NS1, and 1 in NS2. PB2 N701D but not NP T130A affected the temperature dependency of viral transcription. Furthermore, introducing the PB2 701D change into a 5:3 CVV mimic improved and drastically reduced the temperature sensitivity of HA yield. We conclude that RG PR8 backbones used for vaccine manufacture in eggs should contain PB2 701D to maximise virus yield.

## Introduction

Worldwide, annual influenza epidemics result in three to five million cases of severe illness, and 250,000 to 500,000 deaths [1]. Both influenza A viruses (IAV) and influenza B viruses cause seasonal disease but IAV poses additional risks of sporadic zoonotic infections and novel pandemic strains. IAVs are divided into subtypes by their antigenic determinants, the surface glycoproteins haemagglutinin (HA) and neuraminidase (NA). Pandemics have occurred with H1N1 (in 1918 and 2009), H2N2 (1957) and H3N2 (1968) subtype viruses; currently circulating epidemic viruses descended from these are from the H3N2 and 2009 H1N1 (pdm09) lineages.

The primary measure to control influenza is vaccination. Seasonal vaccine production techniques rely on classical reassortment to generate viruses with good growth properties in embryonated hens’ eggs, the major manufacturing substrate. This involves co-infecting eggs with the antigenic (vaccine strain) virus of choice along with a high yielding (“donor”) virus already adapted to growth in eggs. Reassortant viruses that contain the HA and NA of the vaccine viruses are selected and the highest yielding viruses, (high growth reassortants or HGRs), are designated as candidate vaccine viruses (CVVs). Generating HGRs with the desired growth properties can be difficult and sometimes requires further passaging of the initial reassortants to further adapt them to growth in eggs, which can also induce unwanted antigenic changes to the HA [2–7].

An alternative, potentially quicker method to generate HGRs that, conceptually at least, reduces potential antigenic changes, involves using reverse genetics (RG) to create the desired strain [8–10]. This method involves generation of virus by transfection of cells with plasmids encoding the eight genomic segments of IAV which transcribe both viral mRNA and negative sense viral RNA (vRNA), resulting in the *de novo* production of virus particles. Typically, the six viral backbone segments (segments 1, 2, 3, 5, 7 and 8) are derived from the egg-adapted donor strain, whereas the two segments encoding HA and NA are derived from the vaccine strain. This “6:2” reassortant can then be produced in large scale in eggs. RG is the only currently viable method to produce CVVs for highly pathogenic avian IAV strains, since it allows the deletion of polybasic sequences that are determinants for high pathogenicity from the virus HA.

A limited number of donor strains for IAV vaccine manufacture exist. The strain that underpins both classical reassortment and RG approaches is the A/Puerto Rico/8/34 strain (PR8). However, reassortant IAVs with PR8 backbone segments do not always grow sufficiently well to ensure efficient vaccine manufacture, prompting the need for better understanding of the molecular determinants of CVV fitness. Analysis of conventionally derived HGR viruses has shown that as expected, PR8-derived internal segments predominate, with 6:2 and 5:3 (PR8:WT virus) reassortants representing the most common gene constellations. Of the 5:3 HGRs, segment 2 is the most common third vaccine virus-derived segment, especially in human pdm09, but also in H3N2 and H2N2 subtypes [11, 12]. In addition, an avian H5N2 5:3 reassortant was shown to produce higher yields than its 6:2 counterpart [13]. Since all 6 internal PR8 gene segments are presumably adapted to growth in eggs, this preference for the vaccine strain PB1 gene perhaps indicates that it confers a growth advantage in the presence of the vaccine strain HA and/or NA genes. Supporting this, many studies have used RG to confirm that introducing a vaccine virus-derived segment 2 into CVV mimics can improve virus yield for human pdm09 and H3N2 strains, as well as avian H5N1 and H7N9 strains [14–22]. Moreover, it has been shown that CVV 5:3 reassortants containing a pdm09 segment 2 and glycoproteins of avian H5N1 and H7N9 viruses also give higher yields than their respective 5:3 viruses containing the indigenous WT segment 2, suggesting a particular growth advantage conferred to CVVs by the pdm09 segment 2 [22].

The fitness advantage conferred by WT segment 2 may be at the genome packaging level [17, 23, 24], and/or due to a positive contribution from the coding region of segment 2. Segment packaging signals of the glycoprotein genes are known to influence yield [14, 25-32] and it has been demonstrated for H3N2 subtype 5:3 reassortants that the NA and PB1 segments co-segregate, driven by interactions in the coding region of segment 2 [17, 22]. However, this does not exclude contributions from the encoded proteins, complicated by the fact that segment 2 produces at least three polypeptide species: the viral polymerase PB1, a truncated version of PB1, PB1-N40, and from an overlapping reading frame, a virulence factor PB1-F2 [33–35]. Moreover, various PR8 strains are used to make HGRs which can give rise to different growth phenotypes for CVVs containing glycoprotein genes from the same strain/subtype [13, 36]. Overall therefore, a better understanding of the molecular basis for the effects of vaccine strain-derived segment 2s on growth of reassortant IAVs in eggs is needed, to better enable rational design of CVVs.

As a starting point, we rescued CVV mimics that were either 6:2 or 5:3 reassortants between PR8 and pdm09 viruses that differed in whether they contained pdm09 or PR8 segment 2. The expectation, based on empirical evidence and previous studies was that the 5:3 reassortants would grow better than the 6:2 ones. This turned out not to be the case; a result that ultimately led to the identification of PB2 residue 701D as crucial for facilitating the HGR-enhancing characteristics of pdm09 segment 2 in eggs.

## Materials and methods

### Cell lines and viruses

Human embryonic kidney (293T) cells, Madin-Darby canine kidney epithelial cells (MDCK) and MDCK-SIAT1 (stably transfected with the cDNA of human 2,6-sialtransferase; [37] cells were obtained from the Crick Worldwide Influenza Centre, The Francis Crick Institute, London. QT-35 (Japanese quail fibrosarcoma; [38]) cells were obtained from Dr Laurence Tiley, University of Cambridge. Cells were cultured in DMEM (Sigma) containing 10% (v/v) FBS, 100 U/mL penicillin/streptomycin and 100 U/mL GlutaMAX with 1 mg/ml Geneticin as a selection marker for the SIAT cells. IAV infection was carried out in serum-free DMEM containing 100 U/mL penicillin/streptomycin, 100 U/mL GlutaMAX and 0.14% (w/v) BSA. All viruses used in this study were made by RG using previously described plasmids for the PR8 [39], and A(H1N1)pdm2009 strains A/England/195/2009 (Eng195) [40] and A/California/07/2009 (Cal7) [41]. CVV strains NYMC X-179A (X-179A) and NYMC X-181 (X-181) were obtained from the National Institute for Biological Standards and Control (NIBSC) repository. Virus sequence analyses were performed in part using data obtained from the NIAID Influenza Research Database (IRD) [42] through the web site at http://www.fludb.org

### Antisera

Commercially obtained primary antibodies used were: rabbit polyclonal anti-swine H1 HA (Ab91641, AbCam) and mouse monoclonal anti-NP (Ab128193, AbCam). Laboratory-made rabbit polyclonal anti-NP (2915), anti-M1 (2917) and anti-PB2 have already been described [43–45]. Secondary antibodies used for western blot were donkey anti-rabbit DyLight 800 and goat anti-mouse DyLight 680-conjugated (Licor Biosciences). Secondary antibodies used for staining plaque or TCID_50_ assays were goat anti-mouse horseradish peroxidase and goat anti-rabbit horseradish peroxidase (Biorad).

### Site–directed mutagenesis

The QuikChange® Lightning site-directed mutagenesis kit (Stratagene) was used for mutagenesis according to the manufacturer’s instructions. Primers used for site-directed mutagenesis were designed using the primer design tool from Agilent technologies.

### Reverse genetics rescue of viruses

293T cells were transfected with eight pHW2000 plasmids each encoding one of the IAV segments using Lipofectamine 2000 (Invitrogen). Cells were incubated at 37°C, 5% CO_2_ for 6 hours post-transfection before medium was replaced with serum-free virus growth medium. At 2 days post-transfection, 0.5 µg/ml TPCK trypsin was added to cells. Cell culture supernatants were harvested at 3 days post-transfection, clarified and used to infect 10-11 day-old embryonated hens’ eggs (Henry Stewart Ltd). Following incubation for 3 days at 37.5°C, eggs were chilled overnight and virus stocks were harvested, titred and partially sequenced to confirm identity.

### RNA extraction, RT-PCR and sequence analysis

Viral RNA extractions were performed using the QIAamp viral RNA mini kit (QIAGEN) using on-column DNase digestion (QIAGEN). Reverse transcription was performed with the Uni12 primer (AGCAAAAGCAGG) using the Verso® cDNA kit (Thermo Scientific). PCR reactions were performed using Pfu Ultra II fusion 145 HS polymerase (Stratagene) or Taq Polymerase (Invitrogen) according to the manufacturer’s protocol. PCR products were purified for sequencing by Illustra GFX PCR DNA and Gel Band Purification kit (GE Healthcare). Primers and purified DNA were sent to GATC biotech (Lightrun method) for sequencing. Sequences were analysed using the DNAstar software.

### Virus titration

Plaque assays, TCID_50_ assays and HA assays were performed according to standard methods [46]. MDCK or MDCK-SIAT cells were used and infectious foci were visualised by either toluidine blue staining or immunostaining for IAV NP and a tetra-methyl benzidine (TMB) substrate. HA assays were performed in microtitre plates using 1% chicken red blood cells/PBS (TCS Biosciences) and all titres are given per 50 µl.

### Virus purification and analysis

Allantoic fluid was clarified by centrifugation twice at 6,500 × g for 10 mins. Virus was then partially purified by ultracentrifugation at 128,000 × g for 1.5 hours at 4°C through a 30% sucrose cushion. Pellets were resuspended in PBS and in some cases treated with N-glycosidase F (PNGase F; New England Biolabs), according to the manufacturer’s protocol. Virus pellets were lysed in Laemmli’s sample buffer and separated by SDS-PAGE on 10% or 12% polyacrylamide gels under reducing conditions. Protein bands were visualised by Coomassie blue staining (Imperial protein stain, Thermo Scientific) or detected by immunostaining in western blot. Coomassie stained gels were scanned and bands quantified using ImageJ software. Western blots were scanned on a Li-Cor Odyssey Infrared Imaging system v1.2 after staining with the appropriate antibodies and bands were quantified using ImageStudio Lite software (Odyssey).

### Quantitative Real-time PCR

RNA extracted from virus pellets (containing partially purified virus from allantoic fluid pooled from two independent experiments) was reverse transcribed (RT) using the Uni12 primer with the Verso® cDNA kit (Life Technologies), according to the manufacturer’s instructions. qPCR was based on TaqMan chemistry, primers and probes were designed using the Primer express software version 3.0.1 (Applied Biosystems) for Cal7 segments 2 and 6 and PR8 segments 2, 5 and 7. To amplify Cal7 segment 4, Taqman primers/probes were ordered using sequences from the CDC protocol [47]. Due to nucleotide variations between Cal7 and PR8 segment 2, different primers/probe were used to amplify the genes from the two strains. Primer and probe sequences are provided in Table 1. PCR was performed using the Taqman Universal PCR Master Mix (Applied Biosystems), according to the manufacturer’s instructions with the recommended cycling conditions. Samples were run on a QuantStudio 12k Flex machine (Applied Biosystems) and analysed using the QuantStudio 12k Flex software, applying automatic thresholds. Standard curves were generated using serially diluted linearised plasmid containing cDNA of the matching genes or RT products from viruses of known titre. PCR products from both linearised plasmid templates and RT templates were separated on 3% agarose gels, and fragments of the correct size were distinguished. DNA was excised from the gels and extracted using the Illustra GFX PCR DNA and Gel Band Purification Kit (GE Healthcare), according to the manufacturer’s instructions. PCR products were sequence confirmed by Sanger sequencing where sufficient material for sequencing was obtained. qRT–PCR was performed in triplicate per sample and mock-infected-cell, no-RT (with template) and no-template controls both from the RT reaction and for the qRT-PCR mix only were used in each experiment, always giving undetermined C_T_ values for the controls. Relative RT levels were calculated by using C_T_ values for segments from virus pellets from viruses grown at the different temperatures and interpolating from standard curves of RT products of RG 5:3 WT virus grown at 37.5°C for Cal7 segments 2, 4, 6 and PR8 5 and 7 and for PR8 segment 2 from the standard curve of RG 6:2 WT virus grown at 37.5°C.

**TABLE 1.**
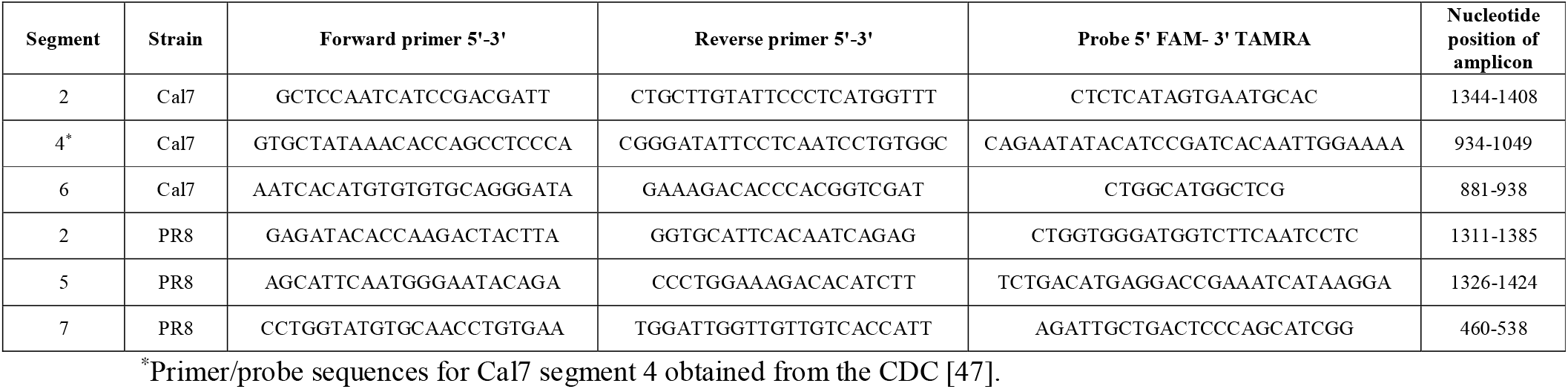
Taqman primers and probes for amplification of Influenza genomic segments by real time RT-PCR.

### Influenza ribonucleoprotein (RNP) reconstitution assays

QT-35 cells at 90% confluency were co-transfected with a chicken pol I firefly luciferase reporter plasmid flanked with segment 8 untranslated regions (UTRs), [48] and four pHW2000 plasmids expressing each of the viral protein components needed to reconstitute RNP complexes using Lipofectamine 2000 (Invitrogen). Triplicate repeats of each assay were performed in parallel at 37.5°C and 35°C. At 48 hours post-transfection, the cells were lysed using Reporter Lysis Buffer (Promega) and luciferase activity measured using Beetle Luciferin (Promega), reconstituted in H_2_O and diluted to a final concentration of 0.6mM. Luciferase activity of each reconstituted RNP was normalised to a ‘No PB2’ negative control.

### Graphs and statistical analyses

All graphs were plotted in and statistical analyses (Tukey’s tests as part of one-way ANOVA) were performed using Graphpad Prism software.

## Results

### Incorporating a pdm09 segment 2 into CVVs confers temperature sensitivity

As a starting point, we used RG to rescue candidate vaccine virus (CVV) mimics that were either 6:2 or 5:3 reassortants between PR8 and the early pdm09 virus isolates Cal7 and Eng195 that differed in whether they contained a pdm09 or PR8 segment 2 in addition to the pdm09 glycoprotein genes. As comparators, parental (non-reassortant) PR8, Cal7 and Eng195 viruses were also rescued. The expectation, based on empirical evidence from existing HGRs as well as from published work that used RG methods [14–22], was that the 5:3 reassortants would grow better than the 6:2 viruses. Viruses were generated by transfecting 293T cells with the desired plasmids, and amplifying virus in eggs. To assess viral growth, TCID_50_ titres were determined on MDCK-SIAT cells. The infectious titre of independently rescued stocks of the 5:3 reassortants were on average ∼ 2-fold higher than the parental pdm09 viruses and ∼ 7-fold higher than the 6:2 reassortants, but around 2 log_10_ lower than WT PR8 (Figure 1A). Surprisingly however, when the HA titres of virus stocks were measured, the PR8/pdm09 6:2 viruses gave on average ∼ 3-fold higher HA titres than the 5:3 viruses (Figure 1B). When HA:infectivity ratios were calculated, the RG 6:2 viruses showed on average ∼ 30-fold higher values than the RG 5:3 viruses (Figure 1C), suggesting an influence of the pdm09 segment 2 on HA content and/or virus particle infectivity.

**FIGURE 1.**
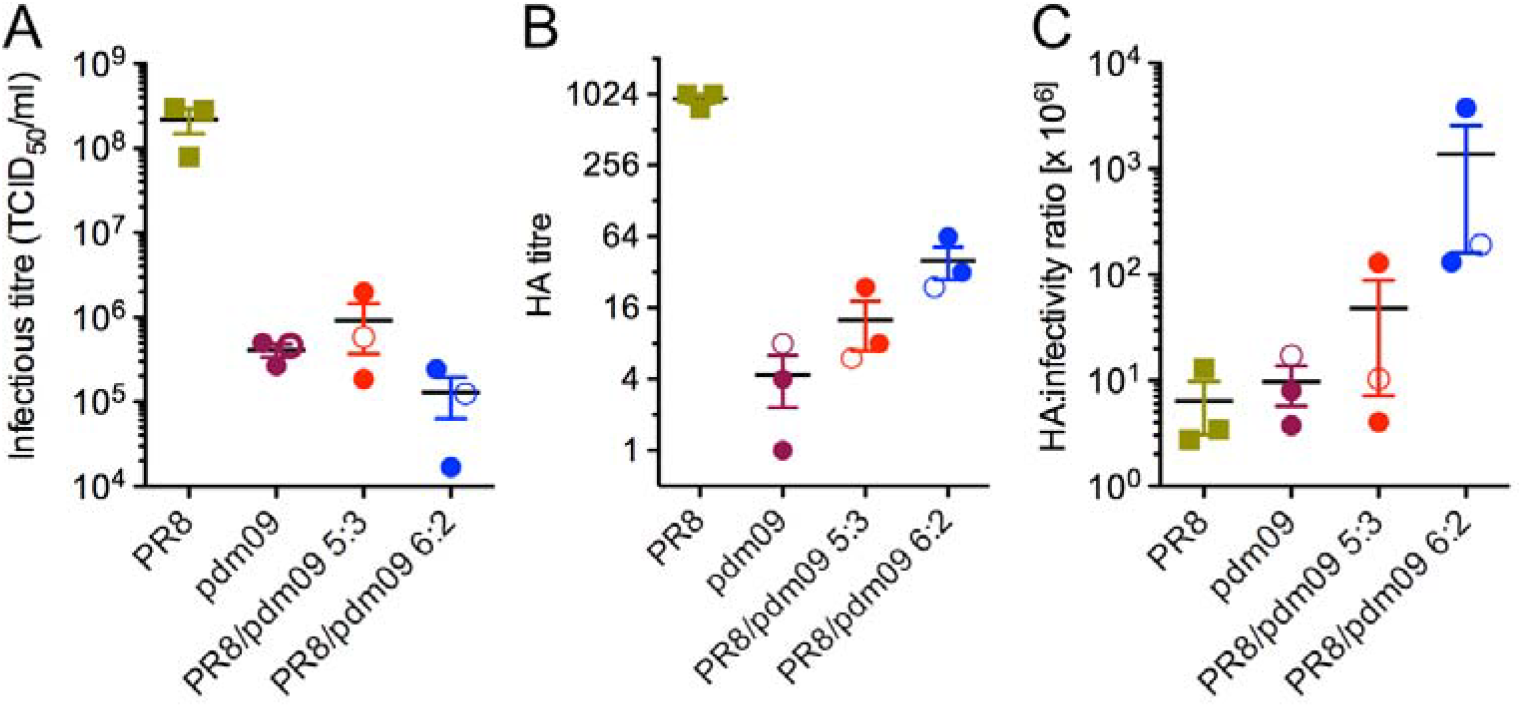
Effect of segment 2 source on virus growth. Virus stocks were grown in eggs and titred by **A)** TCID_50_ assay on MDCK-SIAT cells or **B)** by HA assay. **C)** shows the ratio of HA: infectivity titres, arbitrarily scaled by a factor of 10^6^. Data points are from independently rescued stocks. Filled circles represent viruses with Cal7 glycoproteins and open circles Eng195. Bars represent the mean and SEM.

To further assess the effect of the pdm09 segment 2 on virus yield, eggs were inoculated with a dose range from 10 – 1000 TCID_50_ of virus per egg of the PR8:pdm09 reassortant viruses and allantoic fluid titre measured by HA assay following incubation at 37.5°C for 3 days. The yield of each virus was insensitive to input dose, with no significant differences between average titres within each group of viruses (Figures 2A, B). However, at all doses, the RG Cal7 and Eng195 6:2 reassortants gave higher average HA titres than their 5:3 counterparts, and these differences were mostly statistically significant. As before (Figure 1), this was the opposite of the anticipated result, based on the known compositions of conventionally selected pdm09-based CVVs [11]. However, influenza vaccine manufacture often involves incubation of the eggs at temperatures below 37.5°C [49], so we therefore tested the outcome of growing the reassortant viruses in eggs incubated at 35°C. Again, average HA titres were insensitive to inoculum dose, but the differences between the 5:3 and 6:2 pairs were much reduced and no longer statistically significant. (Figures 2C, D). Growth of both the 6:2 and 5:3 PR8:Cal7 reassortants was improved at 35°C compared to 37.5°C, by around 2-4 fold for the 6:2 virus but by 8-16 fold for the 5:3 virus (Figures 2A, C). Yield of the 6:2 PR8:Eng195 virus was not increased by growth at the lower temperature but substantial gains of around 4-fold were seen with the 5:3 reassortant (Figures 2B, D). Thus the 5:3 viruses including a pdm09 segment 2 appeared to be more temperature sensitive than the RG 6:2 viruses.

**FIGURE 2.**
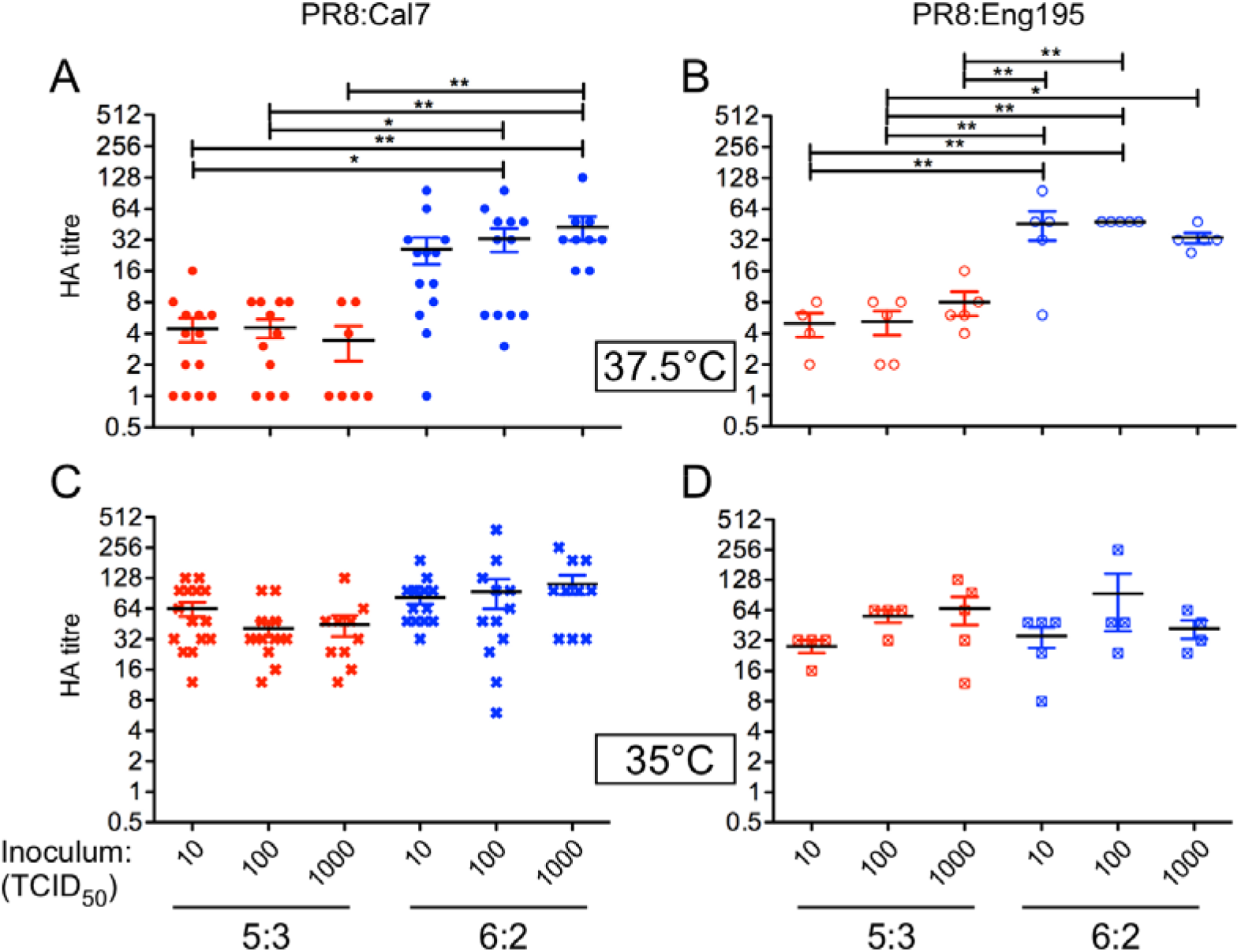
HA yield of PR8:pdm09 5:3 and 6:2 CVV mimics grown at 37.5°C or 35°C. HA titres from allantoic fluid of embryonated eggs infected with reassortants derived from **A, C**) Cal7 or **B, D**) Eng195 grown at 37.5°C (**A, B**) or 35°C (**C, D**) at 3 days p.i.. Bars indicate mean and SEM of 3 independent experiments (5 eggs per condition in an experiment) for PR8:Cal7 reassortants (from two independently rescued RG stocks), and a single experiment for PR8:Eng195 reassortants. Horizontal bars indicate statistical significance (*p < 0.05, **p < 0.01), assessed by Tukey’s test.

### RG 5:3 and 6:2 reassortants differ in their incorporation of HA into virions at different temperatures

To directly assess HA protein yield, viruses from each experiment were partially purified from equal volumes of pooled allantoic fluid by pelleting through 30% sucrose cushions. HA_1_ content from virus pellets was analysed by SDS-PAGE and western blotting either before or after treatment with PNGaseF to remove glycosylation. This gave the expected alternating pattern of slow and faster-migrating HA polypeptide species (Figure 3A, top row). The amount of HA_1_ fluctuated between samples but for both Cal7 and Eng195 reassortants, yield was generally higher from viruses grown at 35°C than 37.5°C and highest from the 6:2 reassortants. To test the reproducibility of this, de-glycosylated HA_1_ was quantified from the western blots of replicate experiments. Absolute HA_1_ yield was variable, but across a total of 5 independent experiments with 4 technical replicates, the average HA_1_ recovery from both PR8:Cal7 and PR8:Eng195 5:3 and 6:2 viruses was improved by growth at 35°C, but by a greater factor (nearly 5-fold versus 3-fold) for the 5:3 reassortants (Figure 3B).

**FIGURE 3.**
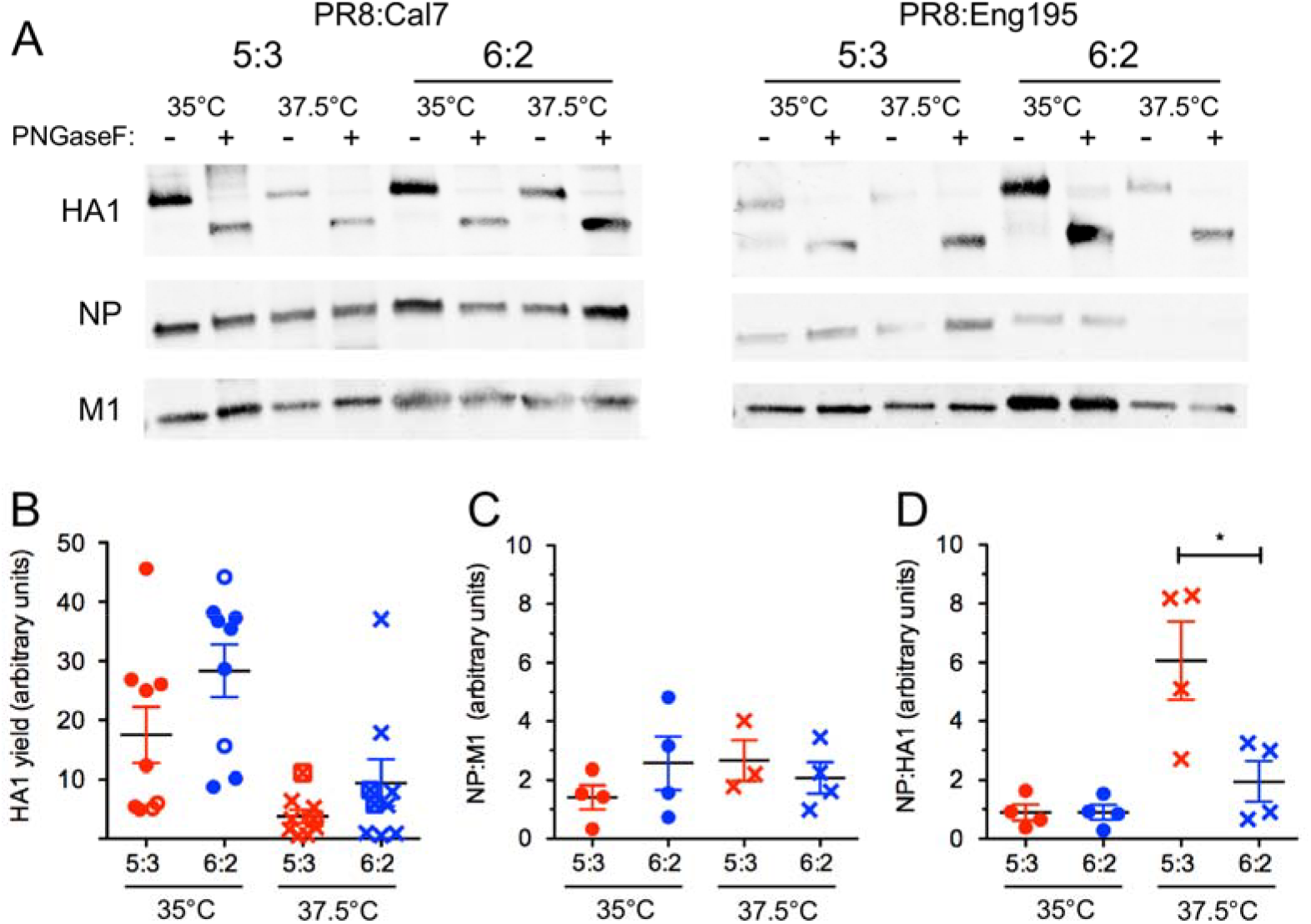
Relative virion composition of viruses grown at 37.5°C versus 35°C. Western blots of purified virus preparations from allantoic fluid of embryonated eggs infected with **A**) PR8:Cal7 or **B**) PR8:Eng195 reassortants grown at 37.5°C or 35°C at 3 days p.i.. Equal volumes of virus samples were either treated with PNGase F (+) or left untreated (-), separated by SDS-PAGE on a 4-20% polyacrylamide gel and virus proteins HA1, NP and M1 detected by western blotting and quantified by densitometry. **C, D, E**) Ratios of NP:HA_1_ (de-glycosylated), NP:M1 and M1:HA_1_ (de-glycosylated) respectively. Bars indicate mean and SEM from 5 independent virus yield experiments (4 experiments with PR8:Cal7 reassortants (filled symbols) using 2 independent RG stocks and a single experiment with PR8:Eng195 reassortants (open symbols)). Horizontal bars indicate statistical significance assessed by Tukey’s test (*p < 0.05).

To test to what extent the varying HA1 yields reflected difference in virus growth and/or HA content of the virus particles, we investigated virion composition by determining the relative amounts of HA_1_ to the other two major structural polypeptides, NP and M1. Western blotting showed reasonably consistent amounts of the latter two proteins in the PR8:Cal7 preparations (Figure 3A, left hand side), but more variable and generally lower recovery of NP in the PR8:Eng195 viruses, especially for the 6:2 virus at 37.5°C (right hand panels). Quantification of these proteins from four independent experiments with the PR8:Cal7 viruses (where the higher growth of the viruses allowed more reliable measurements) showed that the NP:M1 ratios were reasonably consistent and not obviously affected by the incubation temperature of the eggs or the source of segment 2 (Figure 3C). However, the RG 5:3 virus showed a significantly higher NP:HA_1_ ratio than the 6:2 virus when grown at 37.5°C but not at 35°C (Figure 3D). Therefore, the inclusion of the pdm09-derived segment 2 into the PR8 reassortants led to exacerbated temperature sensitivity and lower HA content in virus particles.

### The Cal7 segment 2 does not confer temperature sensitivity to HGRs X-179A and X-181

Following the observation of temperature sensitivity of our RG 5:3 viruses, we tested whether growth of the RG WT pdm09 viruses and corresponding conventional HGR viruses were similarly affected by temperature. Viruses were grown in eggs at 35°C or 37.5°C and the resulting HA titres plotted as fold increases in growth at the lower temperature. Titres of RG viruses containing a PR8 segment 2 were only modestly (∼ 2-4 fold) affected by temperature, but those of viruses containing a pdm09 segment 2 were ∼ 8-16-fold higher at 35°C than 37.5°C (Figure 4; compare solid blue and red bars). However, the yield of the conventionally reassorted authentic 5:3 HGRs X-179A and X-181 (both containing a segment 2 from Cal7 and five other internal gene segments from PR8) were only ∼3-4 fold higher at the lower temperature. Thus, the Cal7 segment 2 gene behaved differently in conventional and RG reassortant virus settings; presumably because of sequence polymorphisms in either segment 2 itself and/or the PR8 backbone between what should be, at first sight, equivalent viruses.

**FIGURE 4.**
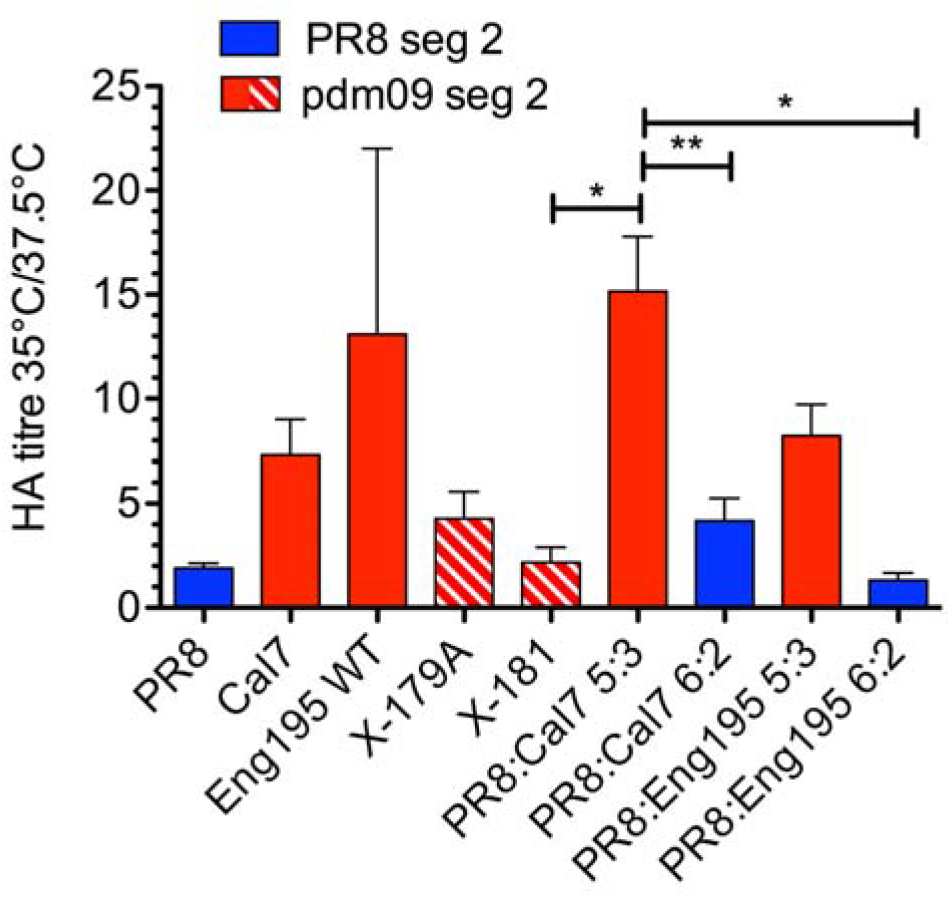
Relative HA titre of RG WT, RG reassortant and HGR viruses containing pdm09 or PR8 segment 2 at 35 °C versus 37.5 °C. For each independent experiment, the fold increase in HA titre of viruses grown at 35°C versus 37.5°C at 3 days p.i. was calculated. Bars indicate mean and SEM from 2-10 independent experiments for each virus. Horizontal bars indicate statistical significance (*p < 0.05, **p < 0.01), assessed by Tukey’s test.

### Internal segments of RG PR8 and HGR X-179A differ

To understand the molecular basis of the temperature sensitivity conferred by RG derived pdm09 segment 2 compared to authentic HGRs, amino-acid sequence comparisons were made between the pdm09-derived genes of the RG viruses used in this study versus those of the HGRs X-179A and X-181. The NA sequences of all four viruses, RG Cal7, RG Eng195, X-179A and X181, were identical (Table 2A). The HA polypeptides of the Cal7, X-179A and X-181 viruses were very similar, differing only with a T209K in the Cal7 sequence and a N129D substitution in the X-181 sequence, while the Eng195 HA varied at four positions from all of the other three viruses and also differed from the HGR viruses in T209K. Within segment 2, the apparent source of the temperature sensitivity, only RG Eng195 differed from the other isolates, with a single amino acid change (R353K). There were no changes in the truncated 11 codon PB1-F2 gene for any of the viruses. Therefore, given the lack of any consistent differences between the two RG pdm09 clones and the conventional HGR viruses, the generally poor and highly temperature sensitive HA yield of the RG 5:3 viruses seemed unlikely to be due to segment 2. Instead, we hypothesised that it was due to epistatic effects arising from sequence differences in the PR8 internal segments of the viruses. Comparison of the internal gene sequences of our RG PR8 and X-179A (no comparable sequences were available for X-181) showed no coding differences in segments 3 and 7, but several in segment 8 (five in NS1 and one in NS2) and one each in PB2 and NP (Table 2B). Amongst these changes, the PB2 N701D polymorphism has been previously linked with host-adaptive changes including temperature sensitivity by several studies [50–58]. Furthermore, PB2 N701D is phenotypically linked with the dominant PB2 host-adaptive polymorphism E627K which also affects temperature sensitive viral polymerase activity [59–61]. This therefore suggested the hypothesis that the PR8 PB2 contributed to the temperature sensitive phenotype seen here.

**TABLE 2.** Amino acid sequence differences between RG CVV mimic and HGR viruses. **A**) Variations from the consensus sequences of the pdm09 PB1, HA and NA polypeptides of the indicated viruses. Sequence accession numbers (segments 2,4 and 6 respectively): Cal7 EPI1355048, EPI1355049, EPI1355051; Eng195 GQ166655.1, GQ166661.1, GQ166659.1; X-179-A CY058517.1, CY058519, CY058521; X-181 GQ906800, GQ906801, GQ906802. **B**) Sequence differences between the backbone-encoded polypeptides of RG PR8 and X-179A. Sequence accession numbers (segments 1,3, 5, 7, 8 respectively): PR8 EF467818, EF467820, EF467822, EF467824, EF467817; X179A CY058516, CY058518, CY058520, CY058522, CY058523.

| Protein | Cal7 | Eng195 | X-179A | X-181 |
| --- | --- | --- | --- | --- |
| PB1 |  | K353R |  |  |
| HA | T209K | L32I,<br>P83S,<br>T209K,<br>R223Q,<br>I321V |  | N129D |
| NA |  |  |  |  |

**B)**
| Protein | No. differences<br>PR8 vs X-179A | Amino acid changes<br>(PR8 > X-179A) |
| --- | --- | --- |
| PB2 | 1 | N701D |
| PA | 0 |  |
| NP | 1 | T130A |
| M1 (M2) | 0 |  |
| NS1<br>(NS2) | 5 (1) | K55E, M104I, G113A,<br>D120G, A132T (E26G) |

To test if the temperature sensitivity conferred by segment 2 of pdm09 viruses could be attributed to effects on viral polymerase activity, we performed RNP reconstitution assays using the readily-transfectable avian QT-35 Japanese quail fibrosarcoma cell line at both 37.5°C and 35°C. Cells were transfected with plasmids to reconstitute RNPs encoding a luciferase reporter gene [59] using either all four PR8 RNP polypeptides, or, to recapitulate RNPs of the 5:3 reassortant virus, PB1 from Cal7 and PB2, PA and NP from PR8. In the latter “5:3” background, the PB2 and NP polymorphisms were tested, singly and in combination, while a negative control lacked a source of PB2. In all cases, increased transcriptional activity of the reconstituted RNPs was observed at the cooler temperature of 35°C, while RNPs containing the Cal7 PB1 protein displayed greater transcriptional activity at both 35°C and 37.5°C than those containing PR8 PB1 (Figure 5A). However, when the ratios of activities at 35°C:37.5°C were calculated, the Cal7 PB1 did not confer greater temperature-dependency on the RNP than the PR8 polypeptide (Figure 5A, green data points). Introducing the PB2 N701D and NP T130A mutations into RNPs incubated at 37.5°C had relatively little effect on viral gene expression, even when both changes were made to reconstitute X179A RNPs. Surprisingly, the PB2 mutation significantly affected RNP activity at 35°C, but by lowering it. Consequently, the ratios of activities at 35°C:37.5°C showed a clear effect of the PB2 (but not the NP) mutation on the temperature-dependency of the RNP. Examination of cell lysates by SDS-PAGE and western blotting for viral proteins PB2 and NP did not show any major differences in their accumulation (Figure 5B). Thus, in the context of a ‘minireplicon’ assay, the Cal7 PB1 did not render RNPs more temperature sensitive, but the PR8 PB2 N701D polymorphism significantly affected the temperature-dependency of the 5:3 virus RNP.

**FIGURE 5.**
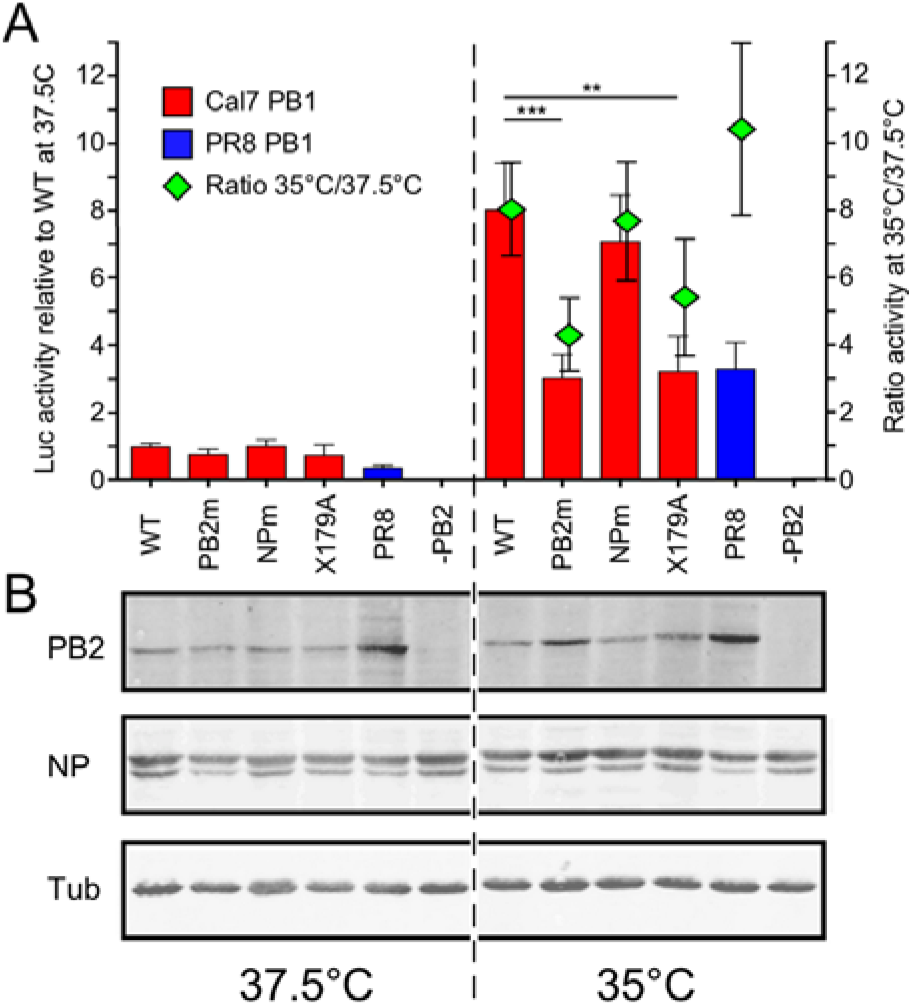
Effect of temperature on RNP activity in avian cells. QT-35 cells were co-transfected with plasmids expressing a synthetic vRNA encoding luciferase along with either Cal7 (red bars) or PR8 PB1 (blue bars) as well as PA, PB2 and NP from PR8, with PB2 and NP either being WT or PB2 N701D (PB2m) and/or NP T130A (NPm) as indicated (RNPs reconstituted with the Cal7 PB1 and both PB2 and NP mutants are equivalent to and labelled as X-179A). Replicate transfections were incubated at 37.5°C or 35°C and at 48 h post-transfection, cells were lysed and luciferase activity measured. **A**) Luciferase activity at each temperature was calculated as fold increases over a negative control lacking PB2 (-PB2), and then normalised to the activity seen from RNPs with Cal7 PB1 and WT PR8 PB2, PA and NP components (WT) at 37.5°C. Data are plotted as bar graphs using the left hand y axis. Statistical significance is indicated (**p < 0.01, ***p < 0.001), assessed by Tukey’s test. To assess the temperature sensitivity of the various RNPs, the ratio of activity at 35°C:37.5°C was calculated and plotted as column means (green diamonds) using the right hand y axis. All values are mean and SEM of 4 independent experiments, with transfections performed in triplicate. **B**) Cell lysates from parallel transfections were analysed by SDS-PAGE and western blotting for viral proteins PB2 and NP. Tubulin (tub) was employed as a loading control.

### PB2 N701D reduces temperature sensitivity of the RG 5:3 virus

To test the significance of the sequence polymorphisms between X-179A and our PR8 internal genes, we attempted rescues of a panel of PR8:Cal7 5:3 viruses using either the WT RG PR8 backbone, PB2 N701D, NP T130A, the NS mutant (NS1 K55E, M104I, G113A, D120G and A132T, and NS2 E26G) or a ‘triple mutant’ containing the mutated PB2, NP and NS genes that would, in protein-coding terms, recreate an RG X-179A. Unexpectedly, viruses with the mutated segment 8 (either singly or as the triple mutant) did not rescue on multiple attempts (data not shown). The reasons for this are not clear, but are suggestive of a detrimental effect on virus replication. However, the PB2 and NP mutants rescued readily and their growth in eggs was further characterised. When HA yield of these viruses at 37.5°C and 35°C was assessed by HA assay, as before the 5:3WT virus was temperature sensitive, giving significantly lower titres at 37.5°C (Figure 6A). The NP mutant behaved similarly to WT at both temperatures, also showing strong temperature sensitivity. In contrast, the PB2 N701D mutant showed a lesser (but still statistically significant) drop in titre at 37.5°C and furthermore, gave significantly higher HA titres than WT at both temperatures. To further test whether the PB2 N701D mutation increased HA yield of the 5:3 CVV mimic, WT and PB2 mutant viruses were partially purified from allantoic fluid and HA content examined by staining with Coomassie blue, with or without prior de-glycosylation. Consistent with the HA titre data, both viruses gave greater amounts of the major structural polypeptides NP, M1 and (best visualised after de-glycosylation), HA following growth at 35°C than 37.5°C, with the PB2 mutant out-performing the WT virus (Figure 6B, upper panel). Levels of de-glycosylated HA_1_ were quantified by densitometry of western blots (Figure 6B, lower panel). Across three replicate experiments, the 5:3WT virus gave on average a 3-fold increase in HA_1_ yield at 35C compared with 37.5°C, whereas the 5:3 PB2 N701D virus showed only a 1.7-fold increase, confirming that the PB2 N701D polymorphism reduced the temperature sensitivity of HA yield in eggs. Finally, we investigated the effects of temperature and the PB2 mutation on the infectivity of the 5:3 viruses. As a measure of virus particle infectivity, we derived genome copy to infectivity ratios for the WT 5:3 reassortant, the PB2 mutant and the authentic X-179A HGR viruses grown at high and low temperatures. RNA from virus pellets was extracted, reverse transcribed and quantitative real-time PCR performed to determine the relative amounts of genome in virions. All viruses incorporated similar levels of segments 2, 4, 5, 6 and 7 and there was no indication of selective defective packaging of a particular segment from any of the viruses grown at the different temperatures (data not shown). Virus infectivity was then determined for each virus sample by TCID_50_ assay and used to calculate genome copy:infectivity ratios, normalised to the virus grown at 35°C. All viruses, including X-179A, showed worse particle:infectivity ratios when grown at 37.5°C (Figure 6C). However, the WT 5:3 RG reassortant virus had an approximately 250-fold higher genome:infectivity ratio than X-179A when grown at 35°C and this was partially (but not completely) restored by the PB2 N701D change. Therefore, having PB2 701D is beneficial to the growth and HA yield of a 5:3 CVV with pdm09 HA, NA and PB1.

**FIGURE 6.**
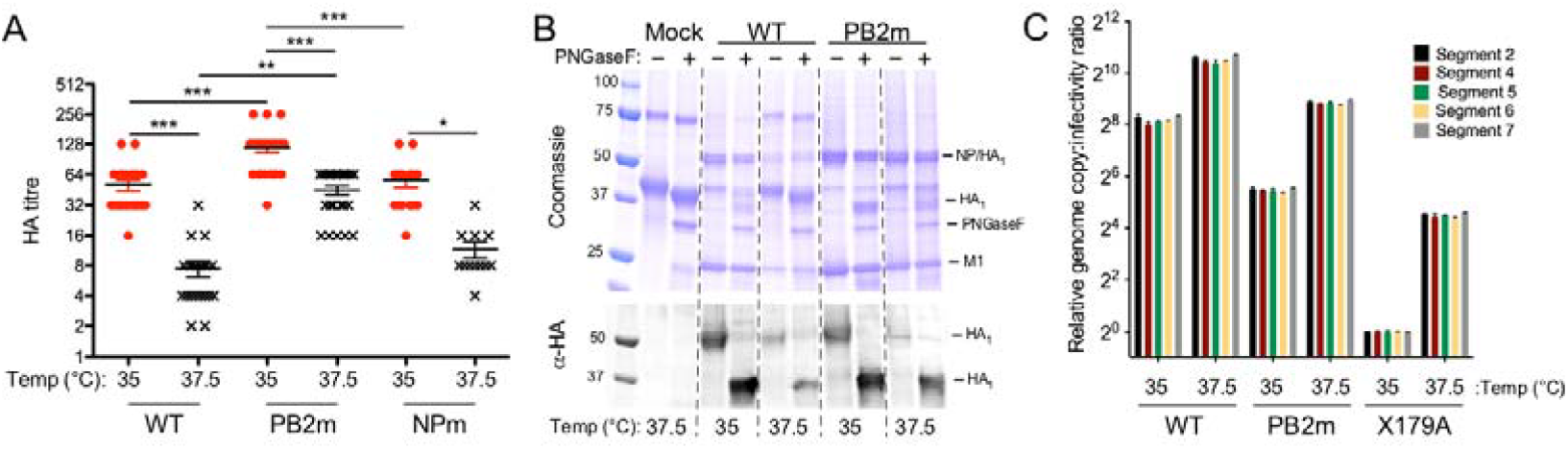
Yield assessment of PR8:Cal7 5:3 mutants grown in eggs at 35°C and 37.5°C. **A**) HA titres from allantoic fluid of embryonated eggs infected with PR8:Cal7 6:2 and 5:3 mutants grown in eggs at 35°C and 37.5°C at 3 days p.i.. Bars indicate mean and SEM from 4 independent experiments using 2 independently rescued stocks of virus for WT and PB2m and 3 independent experiments with a single rescue for NPm (4 - 7 eggs per condition in an experiment). Statistical significance is indicated (*p < 0.05, **p < 0.01, ***p < 0.001), assessed by Tukey’s test. **B**) Representative SDS-PAGE of partially purified virus from pooled allantoic fluid. Equal volumes of each virus sample were either treated with PNGase F (+) or left untreated (-), separated by SDS-PAGE on a 4-20% polyacrylamide gel and analysed by (upper panel) Coomassie staining or (lower panel) western blotting to detect HA1. Molecular mass markers (kDa) are also shown. C) RNA was extracted from virus pellets and qRT-PCR performed to quantify amounts of the indicated segments. Data are plotted as the ratio of genome copy number to infectivity (separately determined by TCID_50_ assay) relative to the value obtained for X-179A grown at 35°C. Error bars reflect the mean and standard deviation of qPCR performed in triplicate per sample.

## Discussion

Several studies in recent years have shown positive effects of incorporating WT segment 2 into RG CVV mimics on yield for human pdm09 and H3N2 strains and avian H5N1 and H7N9 strains [14–22]. In our study, we surprisingly found that for two pdm09 strains, an RG 6:2 virus containing the PR8 segment 2 gave higher HA yield in eggs than the counterpart viruses containing the WT segment 2. Moreover, the RG 5:3 virus had a markedly greater temperature sensitive phenotype compared with the RG 6:2 viruses, as well as with very similar 5:3 genotype classical HGRs. Comparison of amino acid sequence differences between our RG 5:3 viruses and authentic 5:3 HGRs suggested the hypothesis that this was down to epistatic interactions between the WT segment 2 and the internal PR8 genes. Further mutational analysis of the PR8 backbone employed here indicated that the PB2 D701N polymorphism was a major contributor to this genetic incompatibility.

Altering the backbone of our PR8 strain to contain PB2 701D did not completely convert the phenotype of our RG 5:3 CVV mimic to that of its closest authentic HGR counterpart, X-179A in terms of growth in eggs (Figure 6C). It may be that one or more of the other amino acid polymorphisms between the PR8 genes in segments 5 and 8 also contribute. The single difference in NP, T130A, did not affect minireplicon activity (Figure 5) or HA yield in eggs (Figure 6 and data not shown). It lies in the RNA and PB2 binding regions of the protein but the functional significance of differences at this residue are unclear. We were unable to test the significance of the segment 8 polymorphisms as the version of the segment mutated to match that in X-179A could not be rescued into a viable virus, either singly, or when combined with the mutated segment 2 and 5 to supposedly recreate X-179A. The reasons for this are not clear. Possibly by focussing on coding changes only, we missed an essential contribution from a non-coding change (of which there are several between our 5:3 Cal7 reassortant and X-179A, not just in segment 8). Murakami *et al.*, 2008 showed that K55E (in the RNA binding domain) of NS1 mediates growth enhancement of CVVs in MDCK cells [62]. The other amino acid differences are in the effector domain of NS1: position 104 is adjacent to residues known to affect interactions with the cellular cleavage and polyadenylation specificity factor (CPSF), position 113 is in eukaryotic initiation factor 4GI (eIF4GI) binding domain, position 120 is in the 123-127 PKR binding and potential polymerase binding region and position 132 is close to a nuclear export signal (reviewed in [63]). However, any effects of these precise amino acid differences in NS1 and NS2 are not well documented.

The exact mechanism of how PB2 N701D reduces temperature sensitivity of our RG-derived 5:3 virus remains to be elucidated, although our results suggest it may be at the level of viral polymerase activity. Introducing this change into the PR8/Cal7 PB1 polymerase reduced the apparent temperature sensitivity of the viral RNP but by decreasing activity at the lower temperature of 35 °C rather than by increasing activity at the higher temperature (Figure 5). This does not permit a simple correlation to be drawn between the effect of the mutation in the artificial sub-viral minireplicon assay and the behaviour of the complete virus in eggs, but is nonetheless suggestive of a functionally important link. The opposite change, PB2 D701N, has been shown to enhance the interaction of PB2 with mammalian importin α1 [52], so it would be interesting to examine this from the perspective of adaptation to an avian host. Interactions between PB2 and importin α have also been suggested to play a role in viral genome replication [64]; the minireplicon assay used here primarily interrogates transcription, so this could also be an avenue to explore further.

Of the over a hundred PB2 sequences from conventionally reassorted viruses (mostly X-series viruses) available on the Influenza Research Database (accessed December 2018), the vast majority (117/118) have PB2 701D, with a single virus having a glutamate residue. Of the 35 PR8 PB2 sequences available, 701N is a minority variant, only appearing in two viruses; the one used here, and in a “high growth” PR8 derived by serial passage in MDCK cells with the aim of producing a high-yielding backbone constellation for RG vaccine reassortant production in mammalian cells [65]. In this study, the parental PR8 virus possessed PB2 701D before passaging and analysis of reassortant characteristics suggested that this adaptive change was important for growth in cells. Moreover, it has been shown that viruses with PB2 701N were detected in eggs incubated at 33°C but not at 37°C after inoculation with a clinical specimen, suggesting that a lower temperature may be favoured by PB2 701N viruses [66], similar to our study which shows that PB2 701N has a temperature sensitive phenotype. The PR8 clone we used is a descendant of the NIBSC PR8 strain used to make vaccine reassortants, produced by serial passage in MDCK cells [39]; adaptive changes were not determined, but comparison with the NIBSC PR8 PB2 sequence (data not shown) suggests that it did indeed acquire the PB2 D701N change. The data reported here are the reciprocal of those reported by Suzuki and colleagues [65] and further underscore the importance of PB2 701 as a key residue for design of an optimal RG backbone depending on whether the vaccine is to be grown in eggs or mammalian cells.

## Acknowledgements

We thank Dr. Francesco Gubinelli, Dr. Carolyn Nicolson and Dr. Ruth Harvey at the Influenza Resource Centre, National Institute for Biological Standards and Control, U. K. for their support during experiments performed in their lab. We also thank Dr. Helen Wise from the Roslin Institute for useful discussions.

## Funding information

This work was funded in part with Federal funds from the U.S. Department of Health and Human Services, Office of the Assistant Secretary for Preparedness and Response, Biomedical Advanced Research and Development Authority, under Contract No. HHSO100201300005C (to OGE and PD), by a grant from UK Department of Health’s Policy Research Programme (NIBSC Regulatory Science Research Unit), Grant Number 044/0069 (to OGE) as well as Institute Strategic Programme Grants (BB/J01446X/1 and BB/P013740/1) from the Biotechnology and Biological Sciences Research Council (BBSRC) to PD. The views expressed in the publication are those of the author(s) and not necessarily those of the NHS, the Department of Health, ‘arms’ length bodies or other government departments.

SH, MT, RMP and PD declare they have no conflict of interest. OGE and JWMcC have received funding from the International Federation of Pharmaceutical Manufacturers and Associations for the production of influenza candidate vaccine viruses in eggs.

